# *ACTN3* genotype influences skeletal muscle mass regulation and response to dexamethasone

**DOI:** 10.1101/2020.11.20.392282

**Authors:** J.T. Seto, K.N. Roeszler, L.R. Meehan, H.D. Wood, C. Tiong, L. Bek, S.F. Lee, M. Shah, K.G.R. Quinlan, P. Gregorevic, P.J. Houweling, K.N. North

## Abstract

Homozygosity for the common *ACTN3* null polymorphism (*ACTN3* 577X) results in α-actinin-3 deficiency in ~20% of humans worldwide and is linked to reduced sprint and power performance in both elite athletes and the general population. α-Actinin-3 deficiency is also associated with reduced muscle mass and strength, increased risk of sarcopenia in the elderly, and altered response to muscle wasting induced by denervation and immobilisation. *ACTN3* genotype is also a disease modifier for Duchenne muscular dystrophy (DMD), with α-actinin-3 deficiency associated with slower disease progression. Here we show that α-actinin-3 plays a key role in the regulation of protein synthesis and breakdown signalling in skeletal muscle, and its influence on muscle mass begins during early postnatal muscle development. *Actn3* genotype also influences the skeletal muscle response to the glucocorticoid dexamethasone. Following acute dexamethasone exposure, transcriptomic analyses by RT-qPCR and RNA-sequencing show reduced atrophy signalling (*Mstn, Tmem100, mRas, Fbxo32, Trim63*) and anti-inflammatory response in α-actinin-3 deficient mice compared to wild-type. α-Actinin-3 deficiency also protects against muscle wasting following prolonged daily treatment with dexamethasone in female, but not male mice. In combination, these data suggest that ACTN3 R577X is a pharmacogenetic variant influencing the anti-inflammatory and muscle wasting response to glucocorticoid therapy.

## Introduction

Homozygosity for a common null polymorphism in *ACTN3* (R577X) results in complete absence of α-actinin-3 in over 1.5 billion people worldwide (1). α-Actinin-3 deficiency (*ACTN3* 577XX) does not cause muscle disease due to expression of the highly homologous α-actinin-2 in skeletal muscle, but is associated with significantly reduced sprint and muscle power performance in elite athletes and in the general population (2-4). Absence of α-actinin-3 is also associated with reduced skeletal muscle mass that persists through to old age, with an increased risk of sarcopenia, frailty and loss of function in the elderly (5-8). In a case-control analysis of two large independent cohorts of Caucasian post-menopausal women (each *n* > 1200), carriage of the *ACTN3* 577X allele was associated with 33% increased risk of falling (9). A recent study in an elderly Chinese population (age 70-79, *n*=1031) also showed significantly reduced strength and increased frailty score associated with *ACTN3* 577XX genotype (10). *ACTN3* genotype has been shown to modify the clinical severity of various chronic disorders; it contributes to inter-patient variability in disease onset and progression in muscle diseases such as McArdles disease (11), myositis (12), Pompe disease (13) and Duchenne muscular dystrophy (DMD) (14). Moreover, *ACTN3* genotype is correlated with survival in patients with congestive heart failure (CHF); patients carrying the X-allele have 1.72 times higher mortality than patients with *ACTN3* 577RR genotype (*p*=0.01) (15).

Phenotypic analysis of an *Actn3* knockout (KO) mouse model (16, 17) provides some mechanistic explanations for the effect of α-actinin-3 deficiency on skeletal muscle traits and performance. Compared to wildtype (WT) littermates, *Actn3* KO muscles show significant reductions in type 2, fast-twitch glycolytic muscle fibre size, decreased anaerobic activity and increased oxidative phosphorylation (16, 18). This “slowing” of the metabolic properties of fast glycolytic muscle fibres is driven by reduced glycogen phosphorylase activity (19), and is reversed by “rescue” or replacement of α-actinin-3 postnatally in skeletal muscle (20). α-Actinin-3 deficient fast-twitch fibres also show an increase in calcineurin activity (18) as well as alterations in contractile characteristics, with increased rate of decay of twitch transients and changes in calcium signaling caused by increased calcium leak from the SR and reuptake via upregulated expression of SERCA1, calsequestrin and sarcalumenin (21).

It is not yet fully understood how α-actinin-3 deficiency influences muscle mass at baseline and in response to atrophic stimuli. We have previously shown that α-actinin-3 deficiency is associated with reduced muscle wasting and alterations in fibre type switching (compared to WT) in response to hindlimb immobilisation and denervation; this was mediated by elevated baseline calcineurin activity (22). Although calcineurin-dependent pathways are implicated in muscle growth and adaptation to functional overload (23, 24), the α-actinins also directly interact with soluble signalling factors, PI3Kp85 (25), PIP2 (26) and PIP3 (27). All of these factors drive downstream pathways that regulate a number of cellular functions, including the PI3K/Akt/mTOR signalling cascade that regulates protein synthesis and the IGF1-mediated hypertrophic response (28). In cardiomyocyte Z-disks, α-actinin-2 has also been shown to interact with atrogin-1 (*Fbxo32*) an E3-ubiquitin ligase that is upregulated during muscle atrophy (29). In combination, these data suggest that α-actinin-3 deficiency alters muscle mass regulation by modifying key signalling molecules associated with protein synthesis and degradation.

Glucocorticoids are commonly prescribed to treat a variety of conditions associated with chronic inflammation such as Duchenne muscular dystrophy (DMD), but are also known to cause adverse effects such as muscle wasting and osteoporosis (30). In skeletal muscle, glucocorticoids specifically cause atrophy of fast-twitch muscle fibres and elicit muscle wasting by increasing the rate of protein catabolism by the ubiquitin-proteasome and autophagy lysosome system, and by suppressing protein synthesis at the level of translation (31)(32). There is increasing biochemical evidence that members of the α-actinin family play a role in regulating glucocorticoid receptor (GR) activity, raising the possibility that α-actinin-3 deficiency may alter the skeletal muscle response to glucocorticoid treatment (32, 33). α-Actinin-4 (a non-muscle α-actinin) interacts with GR in the presence of the glucocorticoid dexamethasone via the nuclear receptor interacting motif LXXLL, which is conserved across all α-actinin family proteins (34-36) and potentiates GR activity in a dose dependant manner (34-36). Similarly, both α-actinin-4 and α-actinin-2 (which is 80% identical to α-actinin-3 (37)) directly interact with and promote the activity of other nuclear receptors such as the estrogen receptor, androgen receptor and thyroid receptor via the LXXLL motif (34, 36). α-Actinin-2 also binds glucocorticoid receptor interacting protein (GRIP1) and increases its transactivation activities, as well as synergistically enhancing its nuclear receptor coactivator functions (34). These results suggest that deficiency of α-actinin-3, which increases α-actinin-2 expression (38), may likewise alter GR activity and response to glucocorticoids in skeletal muscle.

In this study, we aim to examine the effect of α-actinin-3 deficiency on pathways associated with protein synthesis and degradation in mature skeletal muscle and during development, and then determine if this influences the muscle wasting response to treatment with dexamethasone. We demonstrate that the effect of α-actinin-3 deficiency on muscle mass and downstream mTOR signalling occurs early during postnatal development and prior to full muscle maturation. We further show that α-actinin-3 deficiency reduces the atrophic response induced by dexamethasone, and that α-actinin-3 expression directly influences expression of key genes and pathways associated with muscle adaptation and inflammatory response following dexamethasone treatment. Finally, we demonstrate that α-actinin-3 deficiency protects against muscle wasting induced by dexamethasone in female, but not male mice, and that this is mediated by increased protein synthesis and reduced expression of muscle atrophy-related genes.

## Results

### α-Actinin-3 deficiency alters muscle mass and signalling pathways involved in protein synthesis and breakdown from early postnatal development

Signalling pathways responsible for muscle growth during development, regeneration and overload-induced hypertrophy converge on mTOR and its downstream effectors that control protein synthesis (39). Since the first three weeks of postnatal mouse development is known to be a period of intense growth characterised by an increase in myofibre number (hyperplasia) and myofibre size (hypertrophy) (40), we examined the activation of mTOR via PI3K/Akt, as well as the Smad3, which mediates myostatin/activin A signalling and cross-talks with mTOR signalling via Akt (39), at P0, P7, P14 and P28. We also examined the expression of RCAN1-4 as a marker of calcineurin activity at these developmental time points, since we have previously observed increased calcineurin activity in mature α-actinin-3-deficient skeletal muscle (18). Western blot analysis showed no differences in the activation of these pathways between genotypes at P0, although *Actn3* KO muscles showed significant reductions in p-mTOR and mTOR compared to WT (Supplementary Figure S1). At P7, *Actn3* KO muscles again showed reductions in p-mTOR and overall mTOR activity, as well as reduced 4ebp1 and p-4ebp1; expression of PI3Kp85, Akt, p-Akt, Smad2/3, p-Smad3, S6RP and p-S6RP was not different between genotypes. RCAN1-4 expression was increased in *Actn3* KO muscles compared to WT (Figure 1A, Supplementary Figure S2). In contrast, at P14, mTOR activity and total 4ebp1, but not p-4ebp1, was increased in *Actn3* KO muscles compared to WT; similarly, PI3Kp85 and RCAN1-4 expression were also increased in *Actn3* KO muscles (Figure 1b). Increased activation of Smad3, which is associated with suppression of protein synthesis, was also observed in *Actn3* KO muscles relative to WT at P14. Similar trends were observed at P28 (Supplementary Figure S3) and also in adult, mature skeletal muscles in both male and female mice (Figure 1c, Supplementary Figure S4).

**Figure 1.**
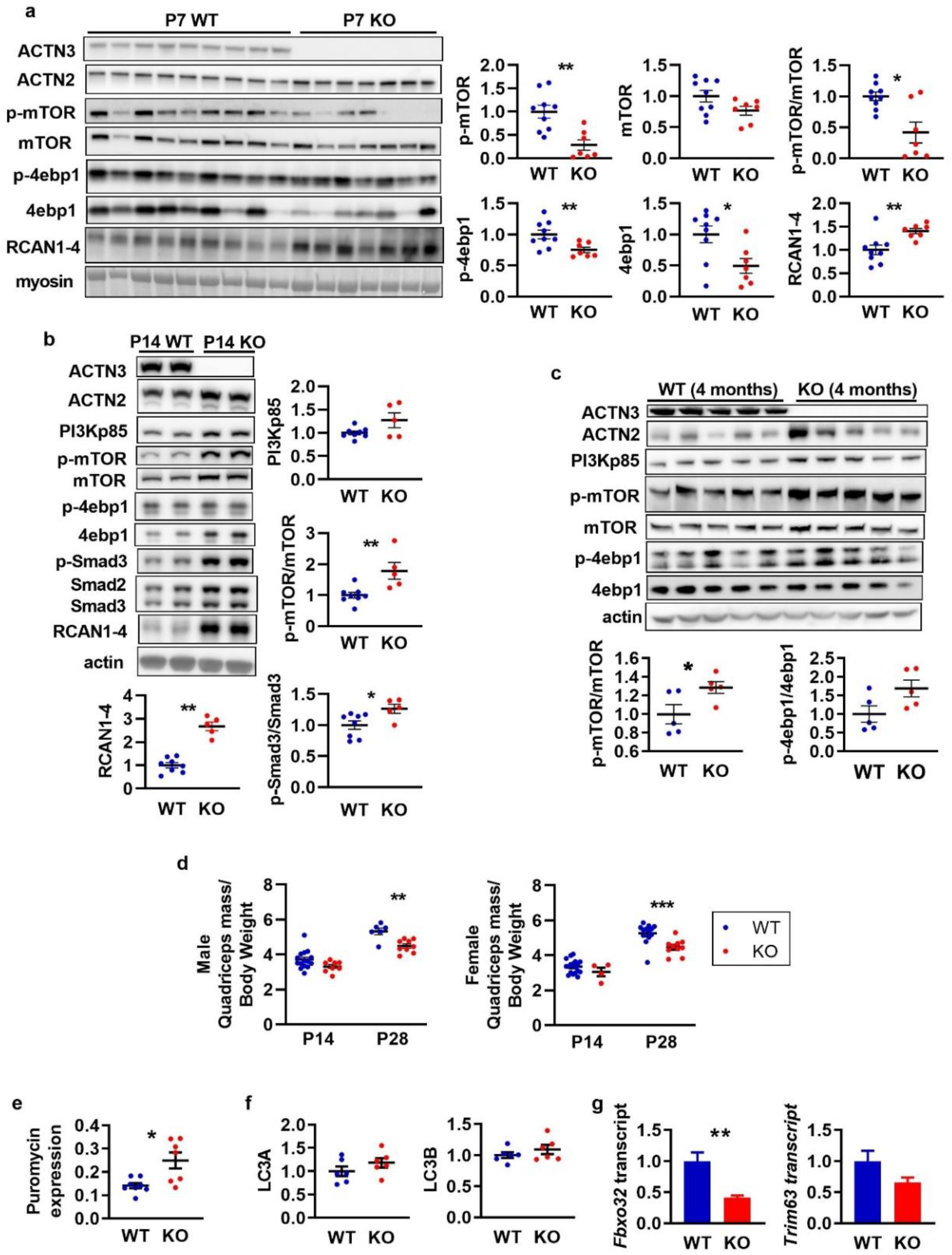
α-Actinin-3 deficiency alters protein synthesis and breakdown signalling in skeletal muscle from early postnatal development. a) At P7, *Actn3* KO quadriceps muscles showed significantly reduced activation of mTOR and 4ebp1 but increased expression of RCAN1-4 compared to WT. (b) At P14, *Actn3* KO muscles showed increased PI3Kp85, RCAN1-4 and activation of mTOR and Smad3 compared to WT. (c) Increased activation of mTOR and 4ebp1 is maintained in adult *Actn3* KO muscles relative to WT. (d) At P28, *Actn3* KO muscles showed significant reductions in quadriceps mass relative to body weight compared to WT. e) Puromycin incorporation (indicator of rate of protein synthesis) is increased significantly higher in male *Actn3* KO muscles compared to WT. (f) Protein expression of autophagy markers LC3A and LC3B are similar between WT and *Actn3* KO muscles, however, (g) transcript expression of E3-ubiquitin ligase genes *Fbxo32* and *Trim63* are significantly reduced in *Actn3* KO. \**P*<0.05, *\*\*P*<0.01, *\*\*\*P*<0.001, Mann Whitney U test. *N* = 4-9 for all experiments.

We have previously shown that α-actinin-3 deficiency is associated with reduced muscle mass in adult mice (16). To determine how changes in protein synthesis with α-actinin-3 deficiency during development correlate temporally with *Actn3* genotype effects on muscle mass, we examined the quadriceps mass of WT and *Actn3* KO mice during development. Muscle mass comparisons between genotypes were performed with muscle mass normalised to body weight to account for differences in litter sizes. Genotype difference in muscle mass relative to body weight was not detected at P0, P7 (data not shown) or at P14, but was significant by P28 in both male and female cohorts (Figure 1D), suggesting that temporal changes in protein synthesis precede the effects of α-actinin-3 deficiency on skeletal muscle mass.

To further examine the impact of these signalling changes associated with α-actinin-3 deficiency on muscle mass regulation in adult skeletal muscle, we compared the rates of protein synthesis between WT and *Actn3* KO muscles *in vivo* using the non-isotopic SUnSET technique (41). Analyses of puromycin incorporation over 30 min demonstrated increased protein synthesis in *Actn3* KO muscles at baseline in males (Figure 1e) although this did not reach significance in females (Supplementary Figure S4). We also examined other markers of autophagy and ubiquitin-proteosome system (UPS) and found no genotype difference in expression of autophagy markers LC3A and LC3B (Figure 1f); however *Actn3* KO muscles show a marked downregulation of *Fbxo32* and *Trim63* compared to WT, which encode for the E3-ubiquitin ligases atrogin-1 and MuRF1, respectively (Figure 1g). Overall, these results suggest that the α-actinin-3 deficiency influences muscle mass and protein synthesis and degradation early during postnatal muscle development, and perturbations of these pathways are maintained in mature adult skeletal muscles in mice.

### *Actn3* KO muscles show reduced dexamethasone-induced muscle atrophy signalling

To determine if the baseline alterations in protein synthesis and breakdown signalling could alter response to muscle wasting in α-actinin-3 deficient muscles, male WT and *Actn3* KO animals were administered with a single pulse of dexamethasone (20 mg/kg) by intraperitoneal injection to activate muscle atrophy signalling and examined at 3 hours and 24 hours post injection. Since α-actinins reportedly influence the activity of glucocorticoid receptor (GR), we first assessed the transcriptional response of upstream GR target genes by RT-qPCR in WT and *Actn3* KO muscles after GR activation. In both WT and *Actn3* KO muscles, treatment with dexamethasone (WT-Dex, KO-Dex) dramatically increased expression of upstream GR target genes *Ddit4, Klf15, and Fkbp5*, as well as *Foxo1 and Foxo3* after 3 hours (Figure 2a). There was no genotype difference in gene expression for the majority of these genes except for *Fkbp5*, where *Actn3* KO muscles showed lower expression relative to WT muscles regardless of treatment. Despite similar upstream response to GR activation, dexamethasone failed to trigger increases in expression for the E3 ubiquitin ligase gene *Fbxo32* in *Actn3* KO-Dex muscles 24 hours post-injection, while WT-Dex muscles showed ~4-fold increase in *Fbxo32* transcript expression relative to WT mice that were given saline (WT-Sal) (Figure 2b). Similarly, WT-Dex muscles show ~2.5-fold increase in *Trim63* transcripts 3 hours and 24 hours post-dexamethasone relative to WT-Sal; while KO-Dex muscles also show 3.6-fold increase in *Trim63* relative to KO mice given saline (KO-Sal) after 3 hours, the increase in *Trim63* transcript expression is not sustained after 24 hours (Figure 2c). Two-way ANOVA showed significant genotype effect on dexamethasone-induced response of *Fbxo32* (*P* = 0.029) and *Trim63* (*P* = 0.007). These data suggest that α-actinin-3 deficient muscles are less responsive to dexamethasone-induced protein degradation. We further examined the expression of major downstream effectors of mTOR that promote protein synthesis. At 24 hours post-dexamethasone injection, WT-Dex muscles showed significant downregulation of p-S6RP, S6RP and p-4ebp1 compared to WT-Sal (Figure 2d), consistent with suppression of protein synthesis signalling. In contrast, KO-Dex muscles showed no significant changes in the expression of these markers relative to KO-Sal (Figure 2e). In combination, these data indicate that α-actinin-3 deficiency results in reduced signalling for muscle atrophy in response to dexamethasone.

**Figure 2.**
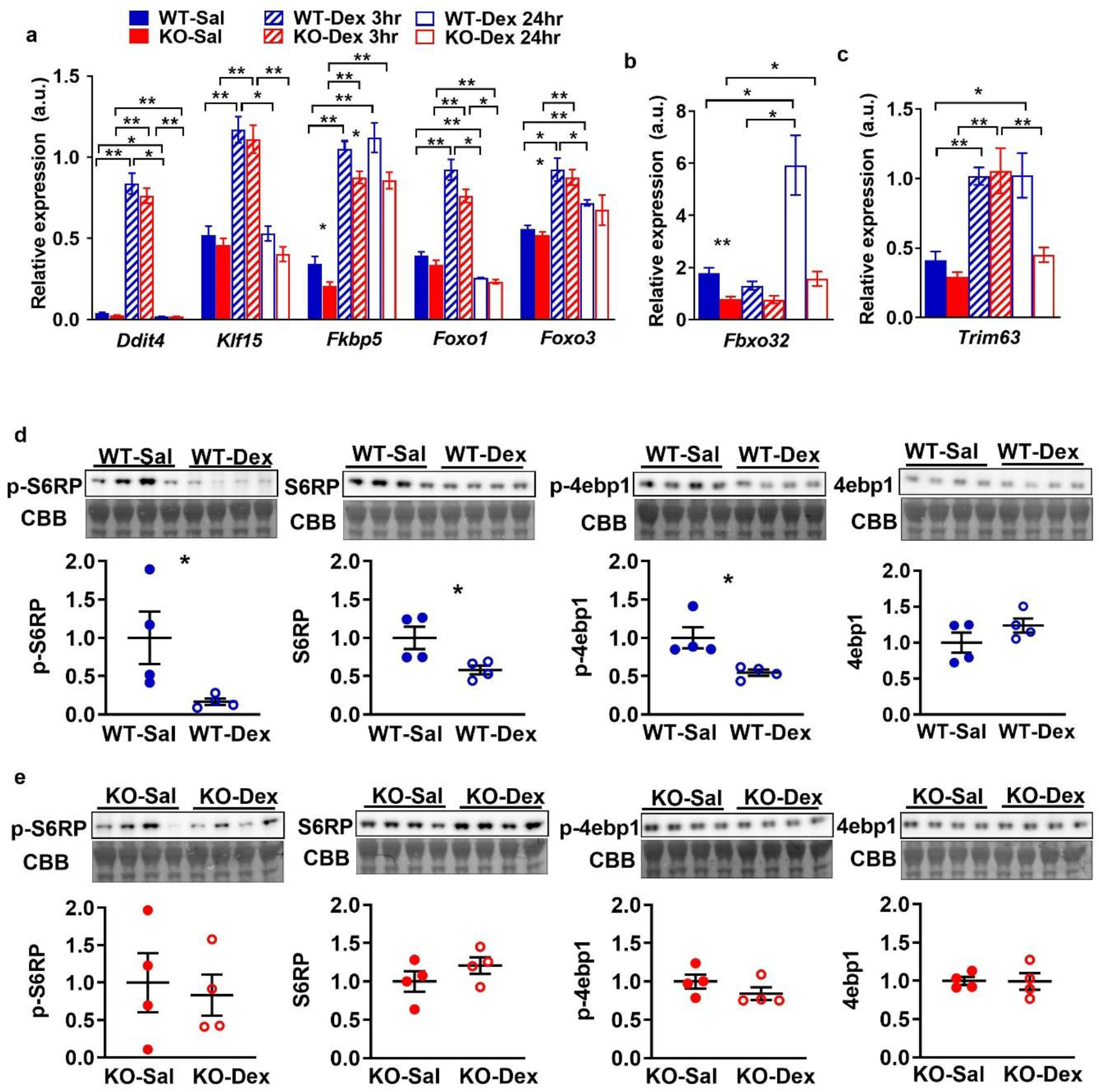
Acute response to dexamethasone in WT and *Actn3* KO mice. a) *Actn3* KO mice show normal transcriptional response to glucocorticoid receptor activation at 3 hr and 24 hr after a single bolus injection of dexamethasone, but fail to activate downstream E3 ubiquitin ligase genes (b) *Fbxo32*, or (c) maintain *Trim63* upregulation at 24 hr post-dexamethasone treatment. (d) Relative to WT-Sal controls, WT-Dex mice show significant downregulation of p-S6RP, total S6RP, p-4ebp1 and no changes to total 4ebp1 at 24 hr post-dexamethasone treatment, consistent with dexamethasone induced suppression of protein synthesis signalling. (e) In contrast, *Actn3* KO-Dex mice are resistant to these changes and show similar levels relative to KO-Sal controls over the same time period. CBB = Coomassie Brilliant Blue. All transcriptional and protein analyses were performed in the quadriceps muscle. \**P*<0.05, \*\**P*<0.01, Mann Whitney U test. *N* = 4-7 for all experiments.

### Global gene expression changes in WT and *Actn3* KO muscles after single bolus dexamethasone injection

We further performed unbiased transcriptomic profiling to determine the acute global gene expression changes in response to dexamethasone injection that are influenced by *Actn3* genotype *in vivo*. Multidimensional scaling analysis of the filtered and processed count data confirmed experimental grouping of gene expression by genotype, treatment and time post dexamethasone treatment (Supplementary Figure S5). Differential expression testing for the effects of treatment, time post-injection, genotype, and genotype interactions with either of the former factors, detected a total of 8653 differentially expressed (DE) genes with *q* < 0.05 across at least one test. Of these, 1416 genes were considered robustly regulated, with both *q* < 0.05 and a fold change greater than 2 for at least one test. Testing confirmed effective transcript-level knockout of *Actn3* expression (Supplementary Figure S6, adjusted *p* =4.44×10^-37^).

The differences in expression of the 1416 robustly regulated genes are presented on a heatmap using genewise centring and scaling, and showed that dexamethasone exhibits its largest influence upon transcription at 3 hours post-injection in both WT and *Actn3* KO (Figure 3a). Volcano plots showed that both dexamethasone treatment (Figure 3b) and *Actn3* genotype (Figure 3c) individually influence transcriptional response. Although correlation plots of dexamethasone response at 3 and 24 hours suggest that WT and *Actn3* KO mice have a broadly similar response to dexamethasone treatment (Figure 3d), specific differential expression testing for interaction effects of *Actn3* genotype on dexamethasone response detected 5 genes of interest, all of which are related to signal transduction pathways associated with muscle growth or wasting, or immune phenotypes: *Tmem100* (*q* = 0.00623), *Cbx8* (*q* = 0.049), *Mras* (*q* = 0.049), *2310022B05Rik* (*q* = 0.0550) and *Mstn* (*q* = 0.1238). Plotting of normalised expression of *Tmem100, Mras* and *Mstn* (which encodes for myostatin) highlight the *Actn3*-mediated difference in expression of these genes in response to dexamethasone (Figure 3e).

**Figure 3.**
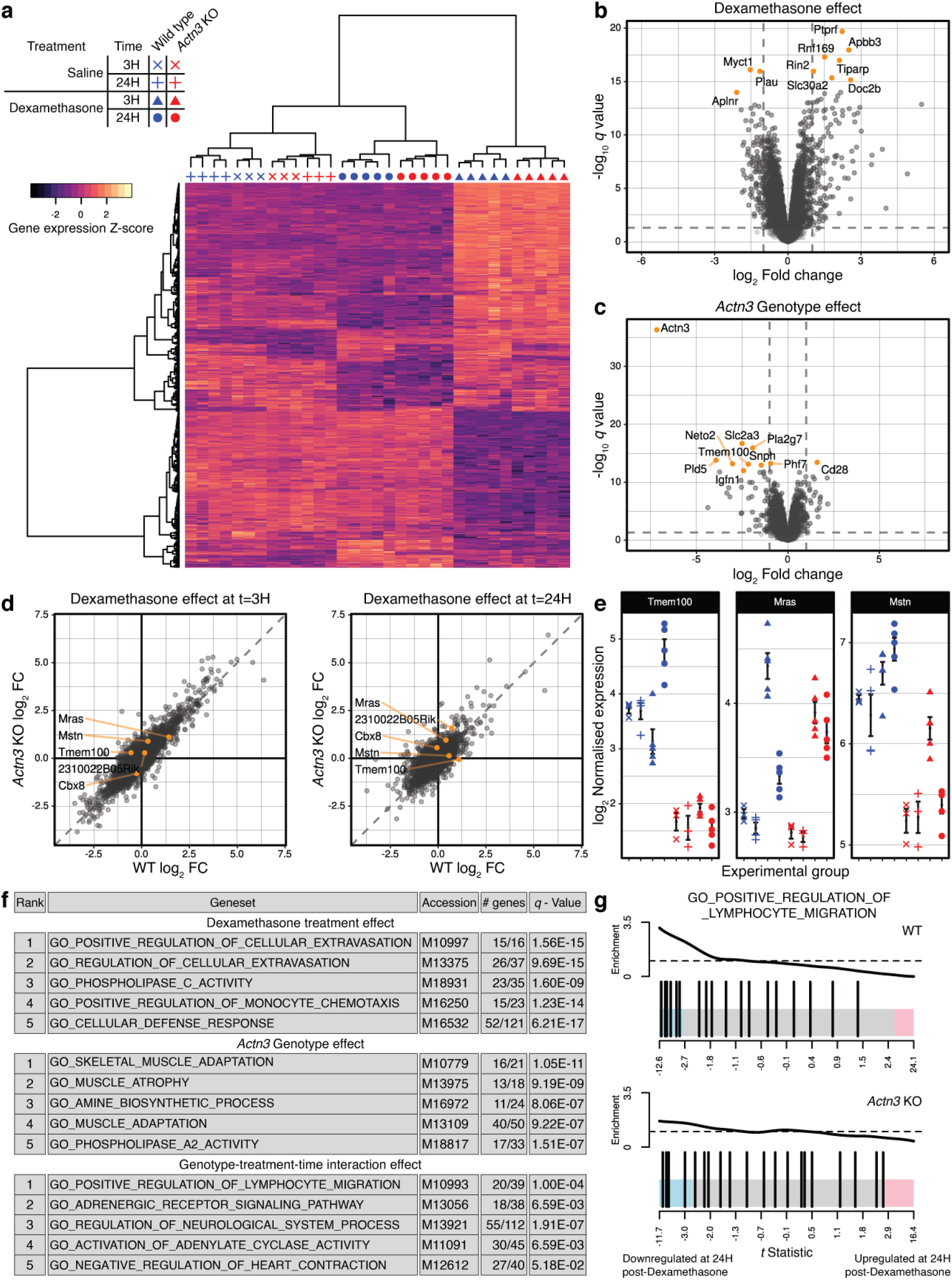
Transcriptomic analysis of WT and *Actn3* KO mice injected with saline or dexamethasone after 3 or 24 hr. An expression heatmap (A) of robustly regulated genes highlights the magnitude of the immediate (3 hr) response to dexamethasone treatment. Expression of robustly regulated genes distinguished experimental conditions, as demonstrated by correlation clustering of samples in the columnar dendrogram. Volcano plots of the effects of dexamethasone treatment (B) and *Actn3* KO (C) demonstrate the efficacy of both interventions and the scale of transcriptional responses. Vertical dashed lines represent a log2 fold change of −1 or 1, and horizontal dashed lines represent a value of *q* = 0.05. (D) Correlation plots show that both *Actn3* KO and WT mice exhibit broadly similar transcriptional responses to dexamethasone at 3 and 24 hours post-injection. Diagonal dashed lines represent y=x, indicating a tendency toward equivalent transcriptional response. Genes with observed q<0.15 for an interaction test between the treatment, time post-injection, and genotype factors are highlighted in orange. (E) The log2 normalised expression of a selection of these genes highlights the difference in dexamethasone response between WT and *Actn3* KO. Symbols represent individual mice, with error bars representing the mean +/- SEM. (F) Gene set enrichment analysis describes a broadly immunosuppressive characterisation of the dexamethasone response and known skeletal muscle phenotypes of *Actn3* KO. All gene sets highlighted for the treatment and genotype effects were associated with downregulation in the dexamethasone-treated and *Actn3* KO animals, respectively. (G) Barcode plots suggest a greater downregulation within GO_POSITIVE_REGULATION_OF_LYMPHOCYTE_ MIGRATION, the top-ranked gene set for the interaction effect, in WT, but not in *Actn3* KO. Enrichment is a relative measure compared to a uniform distribution of *t* statistics within the gene set, represented by the dashed horizontal line.

Gene set enrichment analysis following differential expression testing further characterised the overall response to dexamethasone with respect to *Actn3* genotype (Figure 3f). Analysis showed enrichment of gene ontology terms linked to immune response (“regulation of cellular extravasion”, “regulation of monocyte chemotaxis”, “cellular defence response”) were associated with the dexamethasone effect, while gene ontology terms such as “muscle adaptation” and “muscle atrophy” were associated with the *Actn3* genotype effect. There was also evidence for a differential response to dexamethasone with *Actn3* genotype over time, with gene ontology terms “positive regulation of lymphocyte migration” and “adrenergic receptor signalling pathway” highlighted as ranked by strength of evidence. Barcode plots for “positive regulation of lymphocyte migration” for WT and *Actn3* KO (Figure 3g) indicate suppression of genes related to this pathway in the WT, but not in *Actn3* KO, suggesting that, in the absence of α-actinin-3, skeletal muscles exhibit a diminished immunosuppressive response to dexamethasone.

### Acute dexamethasone treatment increases expression of muscle atrophy-related genes in *Actn3* KO muscles following *ACTN3* replacement

To confirm the role of α-actinin-3 in the altered response to dexamethasone in skeletal muscle, we assessed the dexamethasone induced changes in *Mstn, Tmem100, Fbxo32* and *Trim63* expression in *Actn3* KO muscles following replacement of the *ACTN3* gene (“rescue”). rAAV-CMV-*ACTN3* was injected into the tibialis anterior (TA) muscle of *Actn3* KO mice of one leg and an empty control vector into the TA of the contralateral leg. When stable *ACTN3* expression has been achieved after 4 weeks, mice were further treated with dexamethasone by intraperitoneal injection (20 mg/kg) and TA muscles were harvested after 24 hours (Figure 4a). Comparison of control (−) and corresponding *ACTN3* expressing muscles (+) within each *Actn3* KO mouse by RT-qPCR showed that α-actinin-3 expressing muscles significantly increased transcriptional expression of *Mstn* and *Tmem100* 24 hours after dexamethasone injection (Figure 4b, c), confirming that these genes mediate the altered dexamethasone response associated with α-actinin-3 deficiency. Similarly, expression of *Fbxo32* and *Trim63* were also upregulated following α-actinin-3 replacement in *Actn3* KO mouse muscles (Figure 4d, e).

**Figure 4.**
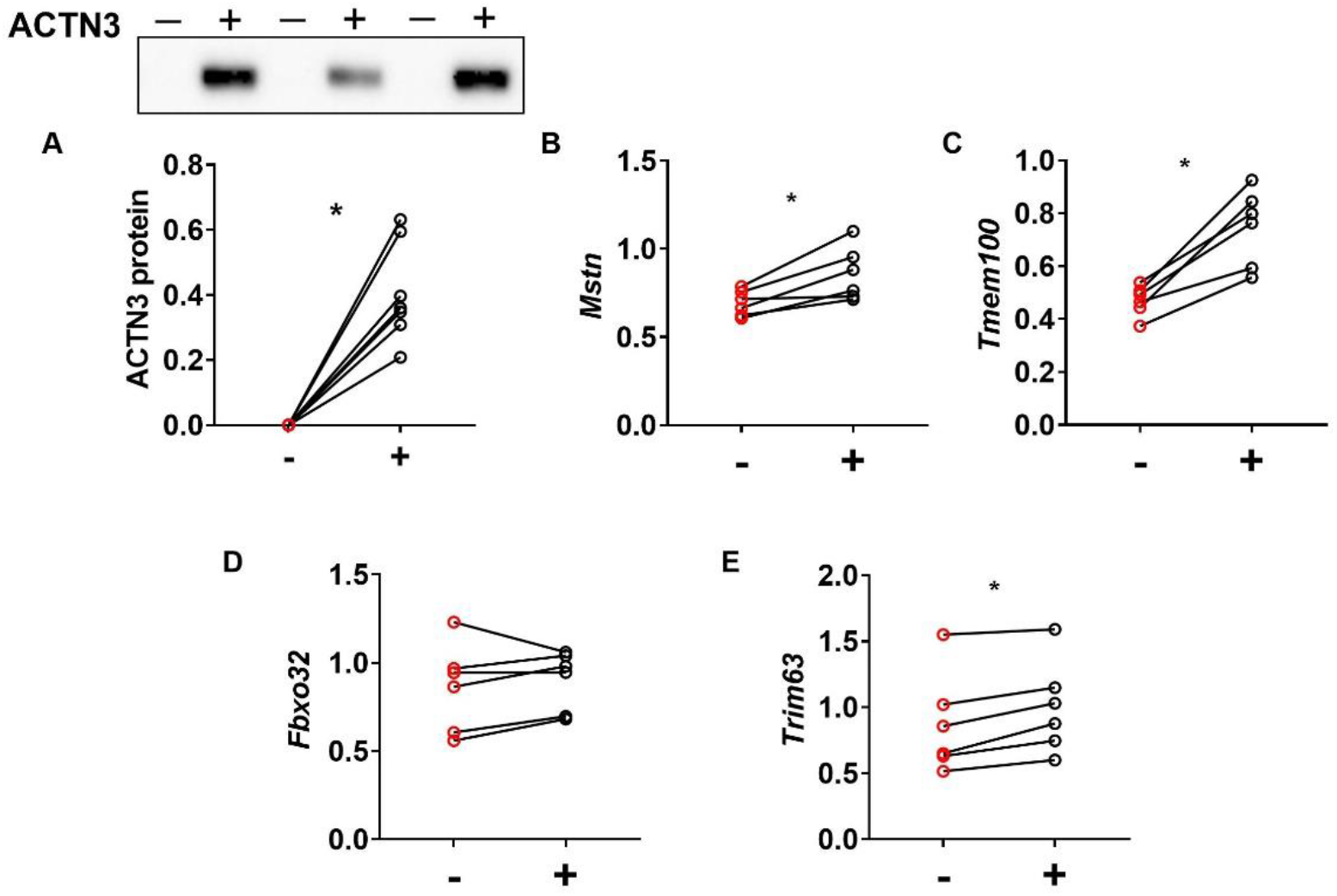
Acute dexamethasone treatment increases expression of muscle atrophy-related genes in *Actn3* KO muscles following *ACTN3* replacement. *Actn3* KO mice were injected with rAAV-CMV-*ACTN3* into the tibialis anterior (TA) muscle on one leg (+) and empty vector (-) into contralateral leg. After 4 weeks of vector expression, mice were given a single dose of dexamethasone at 20 mg/kg. TA muscles were harvested for RT-qPCR analyses after 24 hours. (a) Replacement of *ACTN3* in *Actn3* KO muscles increased expression of (b) *Mstn*, (c) *Tmem100*, (d) *Fbxo32* (in 5 out of 6 mice) and (e) *Trim63*, 24 hours post dexamethasone administration. \**P*<0.05, Wilcoxon test.

### α-Actinin-3 deficiency protects against dexamethasone induced muscle wasting in female mice

To determine if the differential acute response to dexamethasone alters the induction of muscle wasting in α-actinin-3 deficient muscles, WT and *Actn3* KO animals were administered with dexamethasone (20 mg/kg/day) by intraperitoneal injection daily for 2 weeks to induce muscle atrophy. Since glucocorticoids can have sexually dimorphic actions (42), both male and female mice were examined. In males, WT and *Actn3* KO mice showed a similar atrophic response to dexamethasone in the quadriceps muscles; WT-Dex mice showed 11.1% atrophy relative to saline treated mice, while KO-Dex showed 14.1% atrophy (Figure 5a). In contrast, a differential genotype response in muscle atrophy was observed in female mice, with female WT-Dex showing significant muscle atrophy relative to WT-Sal in (−18.6%, *P*=0.0079), while female KO-Dex showed only minimal loss of quadriceps mass relative to KO-Sal (−5.8%, *P*=0.1775). These results were replicated in separate male and female cohorts (Supplementary Figure S7). Two-way ANOVA confirmed the presence of a significant genotype effect on response to dexamethasone for quadriceps mass in females (*P*=0.0045).

**Figure 5.**
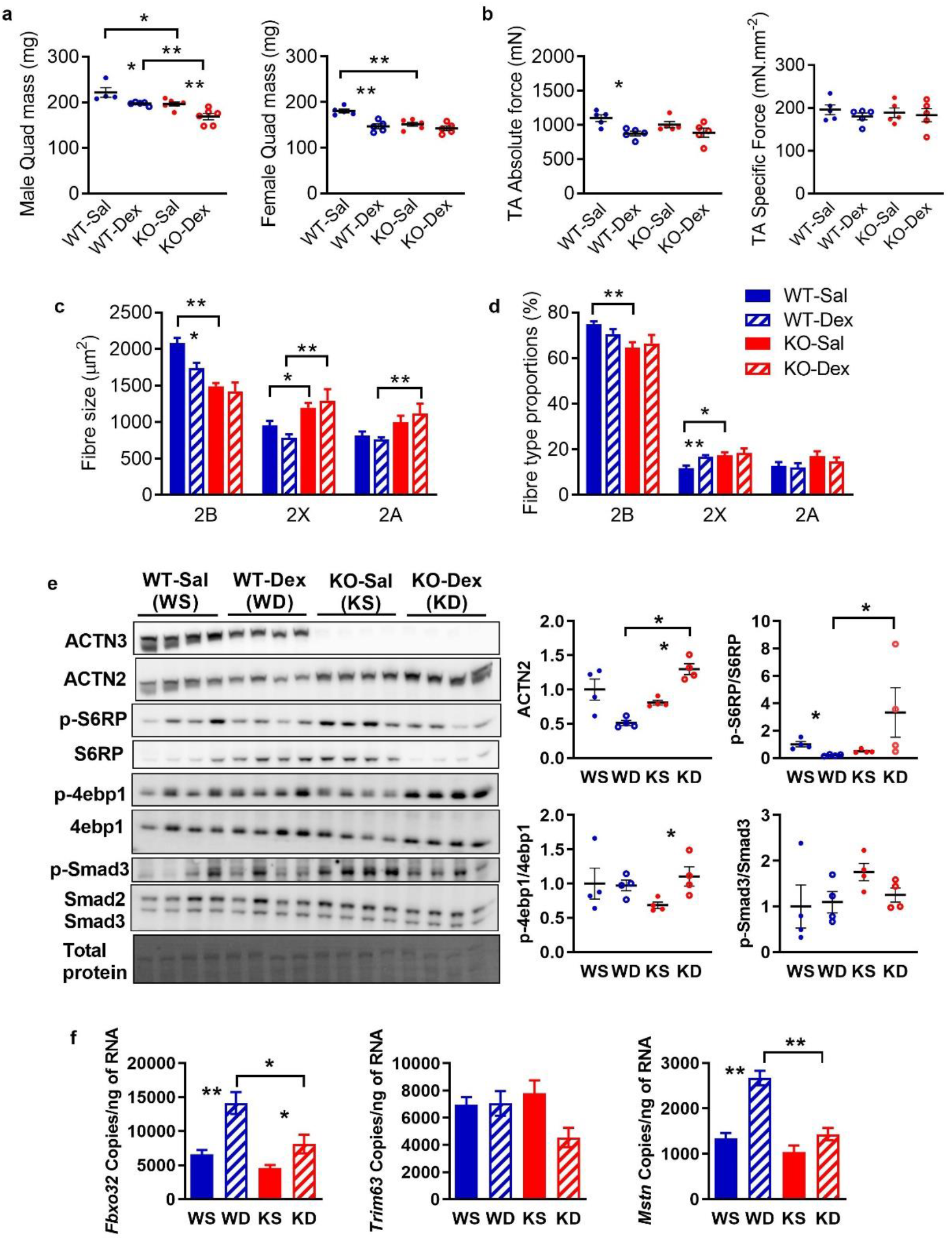
Female *Actn3* KO mice show resistance to muscle wasting induced by prolonged treatment with dexamethasone. A) Male WT and *Actn3* KO mice show similar levels of muscle atrophy in the quadriceps following 2 weeks of daily dexamethasone administration. In contrast, female *Actn3* KO mice showed minimal muscle wasting compared to WT following dexamethasone treatment. B) Female WT-Dex tibialis anterior muscles showed significant decrease in maximal force relative to WT-Sal; specific force is similar regardless of genotype and treatment. C) Female WT-Dex muscles showed reduced fast 2B fibre size compared to WT-Sal as well as a small increase in 2X fibre proportion (D); there was no change in fibre size or proportions in KO-Dex relative to KO-Sal. (E) KO-Dex muscles show increased activation of protein synthesis markers S6RP and 4ebp1, and a trend for decreased Smad3 activity relative to KO-Sal. (F) Transcriptional changes in *Fbxo32, Trim63* and *Mstn* were assessed by ddPCR in WT and *Actn3* KO mice following 7 days of daily dexamethasone injection. WT-Dex muscles showed increased activation of *Fbxo32* and *Mstn* relative to WT-Sal; KO-Dex muscles showed reduced expression of these genes compared to WT-Dex. Scale bar = 50 μm; \**p*<0.05, \*\**p*<0.01 (Mann-Whitney U test). Western blot quantitation values are normalised to total protein and expressed relative to WT-Sal.

Assessment of force generation in the tibialis anterior muscles of female mice also showed significant reductions in WT-Dex relative to WT-Sal (−20.5%, *P*=0.0159), while KO-Dex showed only minimal decrease in maximal force (−7.1%, *P*=0.4286) (Figure 5b). Fibre size analyses of the quadriceps from female mice showed muscle atrophy in response to dexamethasone was restricted to fast 2B fibres in both WT and *Actn3* KO muscles, with the decrease in fast 2B fibre size proportional to the loss in quadriceps mass (WT: −18.9%; *P*=0.0317, KO: −6.6%; *P*=0.5368) (Figure 5c). There was no change in total fibre number (Supplementary Figure S8), but WT-Dex muscles showed a small increase in 2X fibre proportion (1.65%, *P*=0.0079) relative to WT-Sal (Figure 5d).

Markers of protein synthesis signalling (4ebp1, S6RP) and myostatin activation (Smad3) were further analysed (Figure 5e). Compared to WT-Sal, WT-Dex muscles show a small but significant reduction in S6RP phosphorylation, suggesting reduced protein synthesis signalling. In contrast, KO-Dex muscles show marked and significant increases in 4ebp1 phosphorylation, as well as a trend for decreased Smad3 phosphorylation. To further assess transcriptional changes in atrophic signalling following prolonged dexamethasone administration, a separate cohort of female mice were injected with dexamethasone daily for 7 days (Figure 5f). Compared to WT-Sal, WT-Dex muscles showed significant increases in *Fbxo32* and *Mstn* expression, but not *Trim63*, while KO-Dex muscles showed reduced activation of these genes compared to WT-Dex, and a trend for reduced *Trim63* compared to KO-Sal. Significant genotype effect on dexamethasone-induced response was demonstrated for *Mstn* (*P* = 0.004), consistent with decreased dexamethasone induced muscle wasting associated with α-actinin-3 deficiency.

## Discussion

### α-Actinin-3 regulates protein synthesis and breakdown in skeletal muscle from early development

Maintenance of muscle mass requires a balance of protein synthesis and degradation (43). In this study, we demonstrate that loss of α-actinin-3 alters some of the major pathways involved from early postnatal development through to maturity, and thereby regulates skeletal muscle mass during growth and adaptation (Figure 6). The first three weeks of postnatal mouse development is a pivotal period of intense muscle growth (40). Our results show initial decreases in p-mTOR and total mTOR at P0 and P7 and the downstream protein synthesis effector 4ebp1 at P7 in α-actinin-3 deficient muscles. Increased calcineurin activity due to α-actinin-3 deficiency (and α-actinin-2 upregulation) (18) is also detectable from P7. However, from P14, mTOR activation is increased in *Actn3* KO muscles, as is phosphorylation of the transcription factor Smad3, which mediates myostatin-induced atrophic signalling (39). Interestingly, the emergence of this switch in mTOR and Smad3 signalling coincides with the rapid increase in myosin heavy chain 2B expression in mouse skeletal muscle between P7 and P14 (44), and increased myofibrillar protein expression with α-actinin-3 deficiency at P14 (38), suggesting that structural changes associated with α-actinin-3 deficiency may be driven in part by changes in these pathways at this critical juncture in muscle development. The changes seen at P14 persist through to P28 and into maturity, leading to the significant reduction in muscle mass in both male and female α-actinin-3 deficient mice that is detectable from P28.

**Figure 6.**
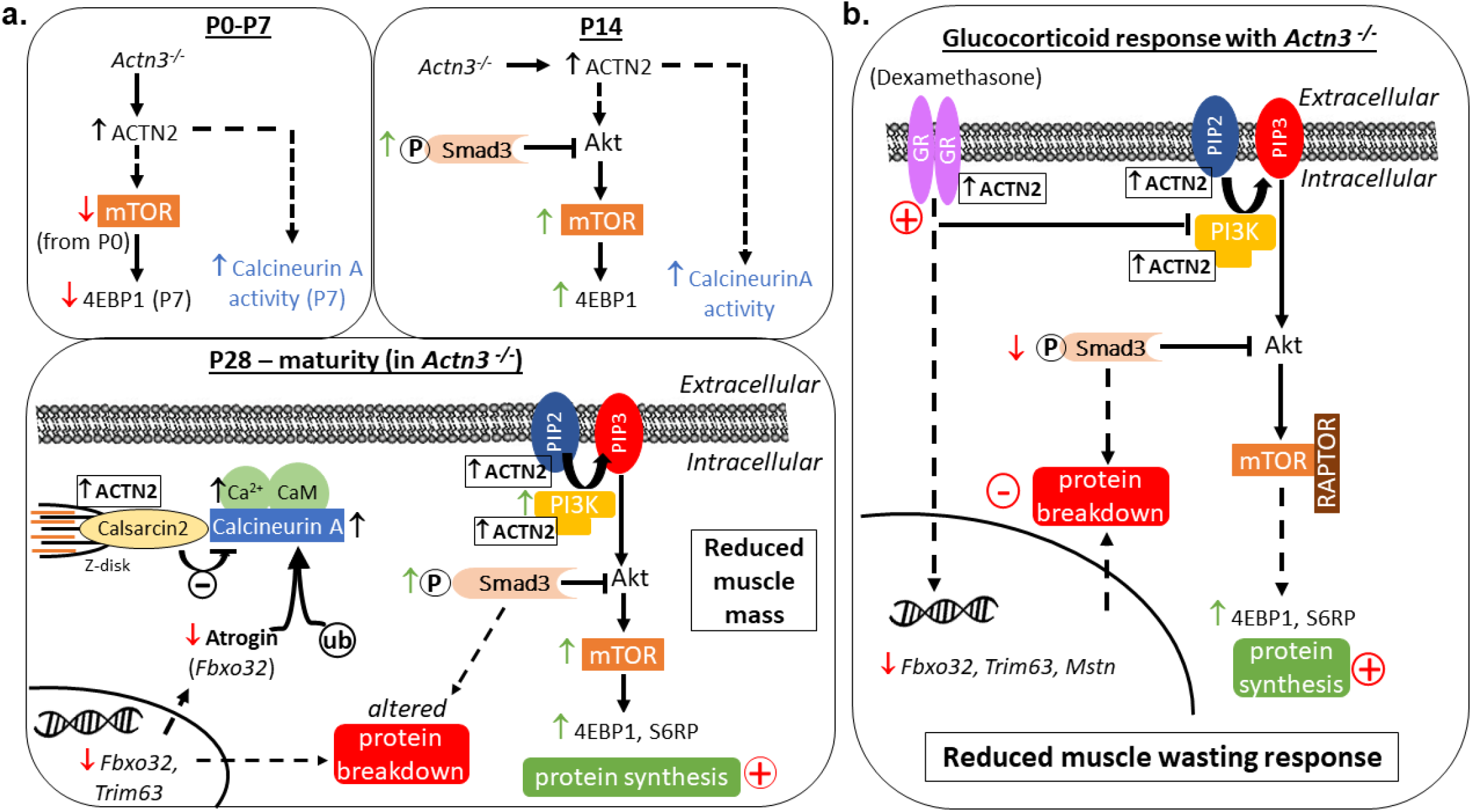
Summary schematic of the effect of α-actinin-3 deficiency on muscle mass regulation and the adaptive response to dexamethasone induced muscle wasting. A) In the absence of α-actinin-3, α-actinin-2 is upregulated to maintain the total sarcomeric α-actinin pool in fast-twitch muscle fibres (38). This is detectable from birth (P0). Decreases in p-mTOR and total mTOR also occur from P0, which subsequently lead to decreases in p-4ebp1 and total 4ebp1 at P7. These are reversed at P14 and increased p-Smad3 is also observed. Increased calcineurin activity (as shown by increased RCAN1-4 expression (18)) occurs from P7. Changes in downstream mTOR and Smad3 signalling in α-actinin-3 deficient muscles persist through to P28 into maturity, resulting in an overall increase in rates of protein synthesis as measured by puromycin incorporation. α-Actinin-3 deficiency also decreases transcriptional expression of E3-ubiquitin ligase genes *Fbxo32* and *Trim63* in muscle. Decreased *Fbxo32* levels with α-actinin-3 deficiency is consistent with increased calcineurin signalling (18) since atrogin-1 targets calcineurin for ubiquitin-mediated proteolysis. These alterations in various protein breakdown pathways along with increased protein synthesis culminate in reduced muscle mass with α-actinin-3 deficiency, which is detectable from P28. B) Despite similar activation of glucocorticoid receptor (GR) by dexamethasone, α-actinin-3 deficient muscles show reduced muscle wasting response due to the inherent baseline effects on protein synthesis and breakdown pathways.

Transcript expression of the E3-ubiquitin ligase genes, atrogin-1 and MuRF1, which promote protein degradation, is also reduced with α-actinin-3 deficiency, consistent with the observed increase in calcineurin activity from P7. In striated muscle, atrogin-1 is known to repress calcineurin activity by targeting calcineurin for ubiquitin-mediated proteolysis, and suppression of atrogin-1 has been shown to increase calcineurin activity in cardiomyocytes (29). Interestingly, α-actinin-2 also directly associates with atrogin-1 at the Z-disk; although α-actinin-2 and calcineurin bind to atrogin-1 on topologically adjacent interaction domains, only calcineurin is targeted for ubiquitination (29), suggesting that α-actinin-2 serves as a dock for recruiting atrogin-1 into the protein complex. In combination, our data suggest that sarcomeric α-actinins play a critical role in regulating muscle protein synthesis and degradation and the homeostatic maintenance of healthy muscle mass from early development, through a combination of their effects on downstream mTOR and Smad3 signalling, reduced atrogin-1 and MuRF1, and regulation of calcineurin activity.

### α-Actinin-3 deficiency attenuates the dexamethasone-induced atrophic response independently of GR activation and protects against muscle wasting

Evaluation of the GR target genes *Ddit4, Klf15* and *Fkbp5* following acute dexamethasone (32) and downstream transcription factors *Foxo1* and *Foxo3*, which act cooperatively with *Klf15* to upregulate atrogin-1 and MuRF-1 (32, 45), indicates that initial GR activation is comparable between genotypes. However, the transactivation response for atrogin-1 and MuRF-1 was different between WT and *Actn3* KO muscles at 24 hr, with α-actinin-3 deficient muscles showing markedly lower expression of both genes. These results suggest that, in the absence of α-actinin-3, muscles are unable to sustain the dexamethasone-induced transcriptional activation of these E3 ubiquitin ligase genes, and that this occurs independently of GR activation. We verified this to be the case by the postnatal replacement of *ACTN3* in *Actn3* KO mice, which led to increased expression of atrogin-1 and MuRF1 24 hr after dexamethasone treatment. In addition, there was no suppression of downstream mTOR protein synthesis effectors in muscles from dexamethasone-treated *Actn3* KO mice, consistent with an attenuated atrophic response. Muscles from female *Actn3* KO mice given prolonged dexamethasone treatment likewise showed reduced atrophic signalling (Smad3, *Fbxo32, Trim63, Mstn)* and enhanced protein synthesis, and were protected from muscle wasting.

### Protection against muscle wasting with α-actinin-3 deficiency is also mediated by the suppression of myostatin signalling

Myostatin is known to mediate dexamethasone-induced muscle atrophy (46), which involves synergistic activation of Akt-FoxO signalling, inhibition of IGF1-PI3K-Akt signalling and increased expression of atrogin-1 mediated by enhanced Smad2/3 signalling (47). Deletion of myostatin prevents muscle atrophy in glucocorticoid-treated mice and blunts induction of proteolytic genes atrogin-1 and MuRF1 (48). In this study, we showed that α-actinin-3 expression is required for the maintenance of myostatin induction. Acute dexamethasone treatment induced increases in *Mstn* in WT muscles that were sustained at 24 hr, while *Mstn* levels returned to basal levels in *Actn3* KO muscles after 24 hrs. Similar results were observed following 7 days of daily treatment. This direct association between α-actinin-3 expression and the maintenance of *Mstn* induction was confirmed by increased *Mstn* levels after 24 hrs following “rescue” in of *Actn3* KO muscle with *ACTN3*. Unbiased transcriptomic profiling of WT and *Actn3* KO muscles following dexamethasone injection showed similar results for *Tmem100*, a gene encoding an intracellular transmembrane protein that is upregulated during activin-A induced muscle wasting (which acts via the myostatin signalling pathway) (49).

### Sexual dimorphic effect of α-actinin-3 deficiency on the response to dexamethasone-induced muscle wasting

The protection against dexamethasone-induced muscle wasting with α-actinin-3 deficiency specifically in female, but not male, mice is intriguing, given that similar changes in baseline protein synthesis and breakdown signalling were observed in both male and female *Actn3* KO mice relative to WT. Dexamethasone has been shown to differentially affect gene expression in the livers of male and female rats (42), and this may be due to cross-talk between GR and other members of the steroid nuclear receptor subfamily, since both estrogen and androgen signalling modulates GR activity in various cell types and tissues (50, 51). α-Actinin-2, which is upregulated in α-actinin-3 deficiency, has been shown to influence androgen and estrogen receptor activities *in vitro* (34). The effects of *ACTN3* R577X on muscle performance in humans also vary with gender; the impact of α-actinin-3 deficiency is more pronounced in elite female athletes compared to males (2). The effect of α-actinin-3 deficiency on sex hormone signalling in skeletal muscle, and how this intersects with atrophic response, requires further study. Gender differences in muscle wasting response have previously been reported in other models. Interestingly, a transgenic mouse model that specifically over-expressed myostatin in skeletal muscle resulted in moderate muscle atrophy (20%) only in males but not females (53)

### ACTN3 genotype influences the anti-inflammatory and muscle wasting response to glucocorticoid

In addition to reduced atrophic signalling, gene set testing demonstrated a diminished immunosuppressive response to dexamethasone with α-actinin-3 deficiency. Two out of five genes of interest that showed significant differential response to dexamethasone between *Actn3* genotypes – *2310022B05Rik* and *Cbx8* (chromobox homolog 8) – are associated with immune phenotypes. In particular, *Cbx8* is a known GR responsive gene. In human lung adenocarcinoma cells, activation of GR with dexamethasone results in downregulation (transrepression) of *Cbx8* through direct binding of GR to its negative glucocorticoid response element (nGRE) (54). Given that GR transrepression activity is generally associated with anti-inflammatory activity and clinical efficacy of glucocorticoids (55, 56), these results suggest that *ACTN3* R577X acts as a pharmacogenetic variant influencing both the anti-inflammatory and muscle wasting response to glucocorticoids in skeletal muscles.

Glucocorticoids are commonly prescribed to treat a variety of chronic degenerative and inflammatory conditions such as Duchenne muscular dystrophy (DMD), but they can cause severe adverse side effects such as weight gain, depression, diabetes, as well as muscle wasting and osteoporosis (30). A recent retrospective analysis on glucocorticoid use in patients with DMD highlighted the variability of its effectiveness and the devastating impact of the side effects on patient well-being, with 12% of patients with DMD in the United States discontinuing therapy due to side effects or insufficient benefit (57). Individual differences in drug response can be due to age, sex, disease, or drug interactions, but genetic factors also play a major role. We have previously shown that *ACTN3* R577X is a genetic modifier of DMD and influences muscle strength and function of young, ambulant patients, as well as disease progression and the age at loss of ambulation (14). Since patients with DMD are typically maintained on glucocorticoids, our results suggest that the association of *ACTN3* R577X with functional outcomes of DMD may be a combination of its effects on both skeletal muscle performance and glucocorticoid response. Our findings confirm *ACTN3* R577X as an important variant to consider in patient stratification for clinical trials of DMD therapeutics to ensure appropriate interpretation of outcomes.

## Conclusion

α-Actinin-3 deficiency is common and occurs in ~1 in 5 people worldwide (1). This study shows for the first time that α-actinin-3 plays a role in the regulation of muscle mass through modulating muscle protein synthesis and breakdown signalling, and that *ACTN3* R577X may influence the glucocorticoid-induced anti-inflammatory and muscle atrophy signalling response. α-Actinin-3 deficiency also protects against dexamethasone-induced muscle wasting, but only in females, suggesting that sex hormone signalling also influences the effect of *ACTN3* genotype on the response to prolonged glucocorticoid use.

## Methodology

### Animals and Ethics

Analyses were performed in muscles from male and female C57BL/6J mice aged P0, P7, P14, P28, 12 weeks and 4-5 months. Experiments involving 2 weeks dexamethasone treatment utilised male and female mice on the C57BL/10ScSn background (aged 4 months, and 10-12 months). Mice were fed standard chow and water *ad-libitum*, and were maintained in a 12:12 hour cycle of light and dark at ambient room temperature (~22°C). All experiments were approved by the Animal Ethics Committee of the Murdoch Children’s Research Institute (MCRI).

### Dexamethasone administration

Mice were administered with dexamethasone at 20 mg/kg/day or saline by intraperitoneal injection either daily for 1 or 2 weeks (chronic response), or with a single bolus injection (acute response; mice were then euthanised at 3 hr or 24 hr post injection). Hindlimb muscles (tibialis anterior, extensor digitorum longus, gastrocnemius, soleus and quadriceps) were harvested and snap frozen in liquid nitrogen or cryopreserved in OCT reagent.

### rAAV

Eight adult male C57BL6 *Actn3* KO mice were anaesthetised (3.5% isoflurane in oxygen) and given temgesic (0.05 mg/kg) for pain management. Hamilton syringes (Hamilton) were used to deliver 30 ul of *rAAV-ACTN3* (20) or rAAV-MCS (control empty vector) diluted in HBSS at a dose of 1E10 vg. Vectors were injected into the anterior compartment of the lower hindlimb targeting the TA and the EDL.

### Muscle Physiology

Force analyses were performed on the tibialis anterior muscles from mice treated with dexamethasone or saline for 2 weeks, using the Aurora Scientific Dual Mode Lever System with supplied software (DMC5 4.5, DMA) and carried out at 37°C as previously described (20).

### Immunoblotting

Snap frozen quadriceps muscles were homogenised in 2% SDS lysis buffer and assessed for total protein concentration using Direct Detect (ThermoFisher Scientific). Proteins were separated by SDS-PAGE using precast midi-criterion gels (Biorad), then transferred to polyvinylidene fluoride membranes (PVDF, Millipore). These were blocked with 5% BSA in 1× TBST, probed overnight at 4°C with primary antibodies against α-actinin-3, α-actinin-2 (ab68204, ab68167, abcam), PI3Kp85 (#4257, CST), Akt1 (#C73H10, CST), p-Akt (Ser463) (#4060, CST), mTOR (#2983, CST), p-mTOR (#5536, CST), p-4ebp1 (#2855, CST), 4ebp1 (#9452, CST), S6RP (#2217, CST), p-S6RP (#4856, CST), Smad2/3 (#8685, CST), p-Smad3 (ab52903, abcam). Blots washed then probed with secondary antibodies at room temperature for 1-2 hours, then developed with ECL reagents (Amersham Biosciences). Membranes were washed, developed and imaged Image Quant (GE Healthcare). Densitometry was performed using ImageJ image processing software (NIH) and quantified using the area under the curve. Some analyses were performed or verified using an automated western blotting technique (*Wes*, Protein Simple). Results were normalized to total protein and presented relative to WT control.

### Surface sensing of translation (SUnSET)

Surface sensing of translation was performed as previously described previously (41). Male and female mice aged 12 weeks were given either a single bolus of 0.04 μmol/g puromycin dihydrochloride/PBS solution (Calbiochem) or PBS alone by intraperitoneal (IP) injection. Animals were euthanised at 30 min post injection. The tibialis anterior muscles were dissected and immediately snap frozen (right leg) and stored at −80°C. To determine the rate of protein synthesis, puromycin incorporation was determined by western blot using a puromycin specific antibody (12D10, Merck).

### Fibre morphometry analysis

Fibre-typing was performed as previously described (58). Sections were imaged on the V-Slide Scanner (MetaSystems) and analysed using Metamorph software (Molecular Devices).

### RNA Extractions and cDNA synthesis

Total RNA was extracted from ~ 50 mg of mouse quadriceps by phenol chloroform extraction (1 mL TRIsure solution; Bioline). RNA was purified using the RNeasy Mini Kit (Qiagen) as per the company’s protocol and eluted in 30μl of Milli-Q water. RNA integrities (RIN value) and total RNA concentrations were then measured using TapeStation (Agilent Technologies 2200) and samples with RINs of 8.5-9.2 were used for further RT-qPCR, DDPCR and RNA-sequencing analyses. Quantification of diluted RNA concentrations was performed using Qubit 3.0 fluorometer (Thermo Fisher Scientific). All RNA samples were further diluted to 25 ng/μl and 1 ng/μl RNA, and 4 μl of 25 ng/μl RNA or 2 μl of 1ng/μl RNA samples and reverse transcribed to synthesise cDNA using the High-Capacity cDNA Reverse Transcription Kit (Thermo Fisher Scientific) as per manufacturer guidelines.

### Droplet Digital Polymerase Chain Reaction (DDPCR) and analyses

Droplet digital polymerase chain reaction assay was conducted using 2X QX200 ddPCR EvaGreen Supermix (Biorad) in a twin.tec 96-well plate (Biorad) to a final volume of 24μl for lipid droplet generation. The plate was heat sealed at 180°C for 5 seconds with foil (PX1 Plate Sealer, BioRad) and centrifuged at 4000rpm for 1 minute before loading into an Auto Droplet Generator (BioRad), where each sample was partitioned into 20,000 uniform sized lipid droplets with the transcripts of interest and cDNA distributed randomly into the droplets. The sample plate of droplets was placed in a thermal cycler (T100, BioRad) for subsequent PCR amplification within each droplet. The thermal cycling conditions were as follows: 1 activation cycle of 5 minutes at 95°C, 40 denaturation cycles of 30 seconds at 96°C and annealing cycles of 1 minute at 55-60°C depending on the target gene of interest, a post-cycling step of signal stabilisation of 5 minutes at 4°C followed by 5 minutes at 90°C. All cycling steps were performed using a 2°C per second ramp rate. Following PCR amplification, the sample plate was loaded on the QX200 Droplet reader (Biorad) and the assay information was entered into the software QuantaSoft (BioRad). Droplets were analysed by the droplet reader in the Evagreen fluorescent channel. The fraction of positive droplets in a sample were counted and fitted to a statistical Poisson algorithm to provide plot readouts of absolute quantification and concentration of the transcript of interest. Samples with lipid droplet counts less than 10,000 were rejected. The 1D amplitude plot of fluorescence amplitude was used to determine data quality based on positive and negative droplets separation, droplet scattering and channel amplitude signals (minimum of ~5400). A baseline threshold was applied equally across all samples depending on the fluorescence amplitude to distinguish positive from negative droplets. Readings of the target were adjusted for RNA loading and presented as absolute values of RNA transcript copies/ng of RNA.

### RT-qPCR

cDNA was generated using 500 ng of total RNA using the High-Capacity cDNA Reverse Transcription Kit (Thermo Fisher Scientific) as per manufacturer guidelines. RT-qPCR was performed in triplicate with the LightCycler 480 instrument (Roche Diagnostics) and LightCycler 384-well plates with sealing foil (Roche). Reaction volume of 10 μl contains 2x SensiFAST SYBR No-ROX mix (Bioline), 0.4 μl of each 10 μM forward and reverse primers and 1 μl of 1:10 diluted cDNA. Amplification of single PCR product was confirmed using the melting point dissociation curve and *Cp* values were calculated using the LightCycler 480 software. For each gene, a standard curve was generated using a WT-Sal sample (sample cDNA neat, 1 in 5, 1 in 10 and 1 in 100) and used to convert *Cp* values to relative gene expression for all unknown samples. Gene expression is further normalised to the geomean of three housekeepers *Aldoa, Rer1, Rpl7L1* (59). Primers for qPCR and ddPCR reactions are as follows: *Tmem100* (forward: CTTTCCCAGAAGTTGAACG, reverse TGCAAGCTCACAGAAAGG), *Klf15* (forward: CCCTTTGCCTGCACCTGG, reverse: TGGTACGGCTTCACACCC), *Ddit4* (forward: CTGGAGAGCTCGGACTGC, reverse: CCCATCCAGGTATGAGGAG), *Fkbp5* (forward: GCCATCGTGAAAGAGAAGGG, reverse: CAGCCAGGACACTATCTTCC), *Foxo1* (forward: ACGAGTGGATGGTGAAGAGC, reverse: GGACAGATTGTGGCGAATTG), *Foxo3* (forward: ATAAGGGCGACAGCAACAGC, reverse: CGTGCCTTCATTCTGAACG)*, Fbx032* (forward: ATTCTACACTGGCAGCAGCA, reverse: TCAGCCTCTGCATGATGTTC)*, Trim63* (forward: ACCTGCTGGTGGAAAACATC, reverse: AGGAGCAAGTAGGCACCTCA), *Mstn* (forward: GTTCATGCTGATTGCTGCTG, reverse: CACGCACATGCATTACACAG)*, Actn3* (forward: TTCAACCACTTTGACCGGAA, reverse: CACCATGGTCATGATTCGAG), *Aldoa* (forward: ACATTGCTGAAGCCCAACAT, reverse: ACAGGAAAGTGACCCCAGTG)*, Rer1* (forward: GCCTTGGGAATTTACCACCT, reverse: CTTCGAATGAAGGGACGAAA)*, Rpl7L1* (forward: ACGGTGGAGCCTTATGTGAC, reverse: TCCGTCAGAGGGACTGTCTT).

### RNA-seq analysis

RNA extracted from quadriceps tissue from a total of 17 WT and 16 KO male mice across treatment groups (saline/Dex, at t=3h or t=24h) underwent RNA sequencing using the Illumina HiSeq 2500 sequencing system per manufacturer instructions. Raw read data were processed using the Illumina BaseSpace RNA Express application (60) per standard operating instructions. Reads were aligned using STAR ultrafast RNA seq aligner (61) in the SAM file format (62), then counted using HTSeq (63). Analysis of the resultant genewise count data was performed using the R statistical programming language (version 3.5.0) (64). The dataset was filtered for genes registering more than 0.5 counts per million in at least 2 samples. This resulted in retaining 13,097 genes for analysis. Count data was then normalised and processed for further analyses using the voom function (65) in the *Limma* software package version 3.36.2 (66). Normalised gene expressions were fit to a linear model and differential expression was tested using empirical Bayes methods (67). For the purposes of differential expression testing, samples taken from animals injected with saline at t=3h and t=24h post injection were treated as a single saline-injected grouping. Multidimensional scaling plots and hierarchical cluster analysis both failed to detect a robust difference between these time points, validating the decision to pool sample groups for increased statistical power. The false discovery rate of multiple comparisons was controlled using Benjamini and Hochberg’s correction (68). Genes with FDR<0.05 and fold change >2 for any contrast were considered robustly regulated and hierarchically clustered using a correlation distance metric. Expression data were characterised using the fast pre-ranked implementation (69) of the gene set enrichment analysis (GSEA) method (70). Gene set tests were conducted using the mouse orthologue conversions (71) of Hallmark (72) and C2 curated gene sets from the molecular signatures database (MSigDB 3.0) (73).

### Statistics

Analyses were performed in Graphpad Prism (V7, Graphpad Software Inc.) and StataSE (StataCorp). As group sizes consisted of <12, two-sided, unpaired t-tests using nonparametric statistics, Mann-Whitney U-tests were applied using an alpha of 0.05 for all analyses involving independent samples and Wilcoxon signed-rank test for dependent samples. Data were presented as individual points with the calculated group mean (line), or as bar graphs, with ±SEM error bars for each group.

## Acknowledgements

We thank Dr Susan Donath for her assistance with statistical analyses. This work is supported by the Australian National Health and Medical Research Council (NHMRC) project grant (APP1130215) awarded to JTS, PG and KNN. MS is supported by an Australian Government Research Training Program (RTP) Scholarship. KGRQ is a Scientia Senior Lecturer at UNSW Sydney.

