## Supplementary Figure S1 for "*ACTN3* genotype influences skeletal muscle mass regulation and response to dexamethasone"

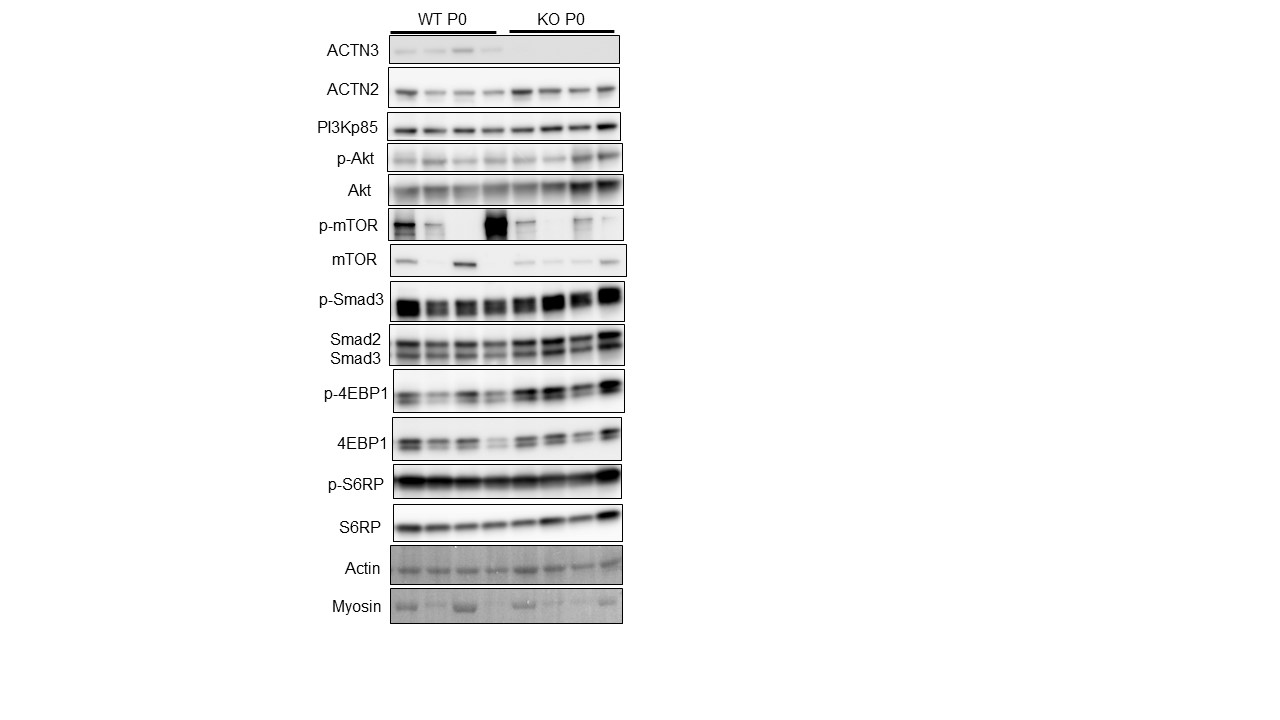

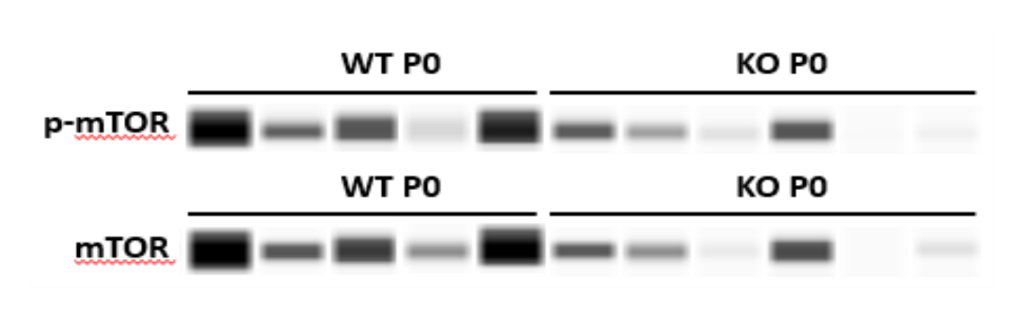

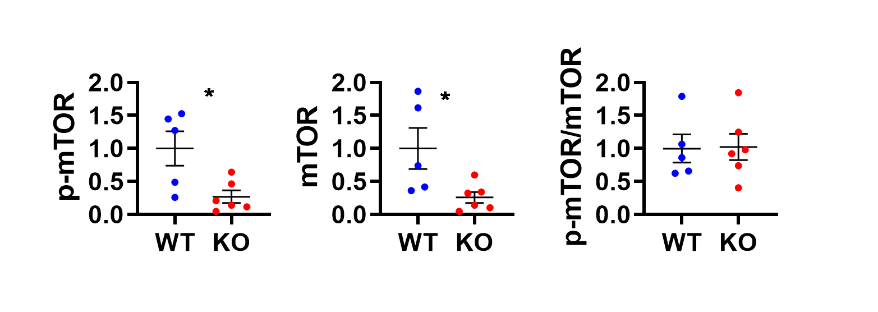

**Supplementary Figure S1.** No difference in expression of α-actinin-2, PI3K/Akt/mTOR, activation of downstream mTOR protein synthesis markers 4EBP1 and S6RP or Smad3 and between WT and *Actn3* KO muscles at P0. However, p-mTOR and total mTOR are significantly downregulated in expression in KO muscles compared to WT in a repeat experiment using automated western blotting.
