## Supplementary Figure S8 for "*ACTN3* genotype influences skeletal muscle mass regulation and response to dexamethasone"

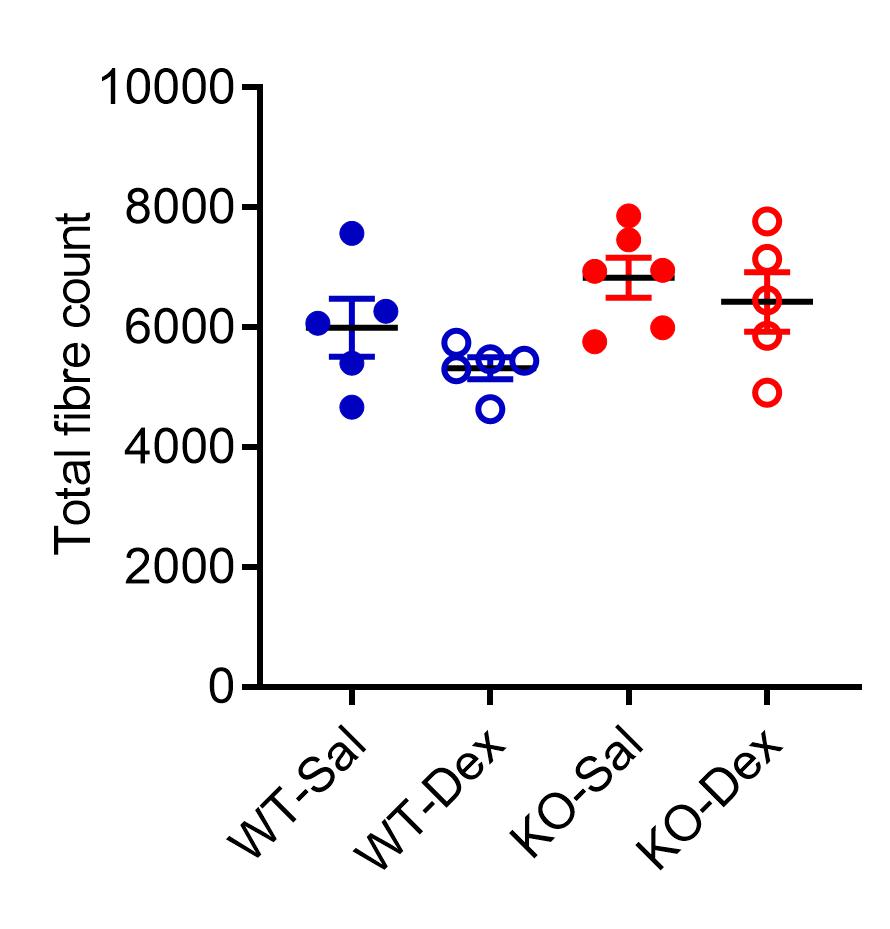

**Supplementary Figure S8.** No significant changes in total fibre number between *Actn3* genotypes or with dexamethasone treatment.
