## Supplementary Figure S2 for "*ACTN3* genotype influences skeletal muscle mass regulation and response to dexamethasone"

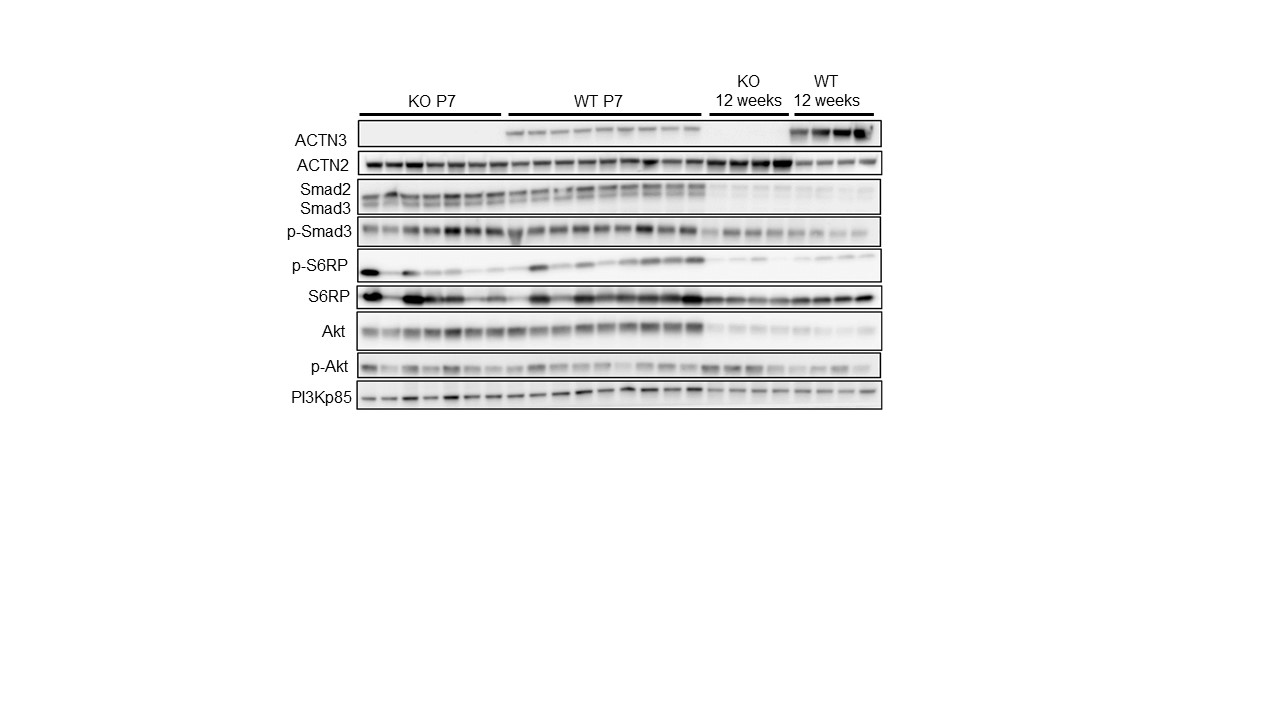

**Supplementary Figure S2**. Expression of PI3Kp85, Akt, p-Akt, Smad2/3, p-Smad3, S6RP and p-S6RP was not different between genotypes at P7. ACTN2 is upregulated in *Actn3* KO muscles at P7.
