## Supplementary Figure S3 for "*ACTN3* genotype influences skeletal muscle mass regulation and response to dexamethasone"

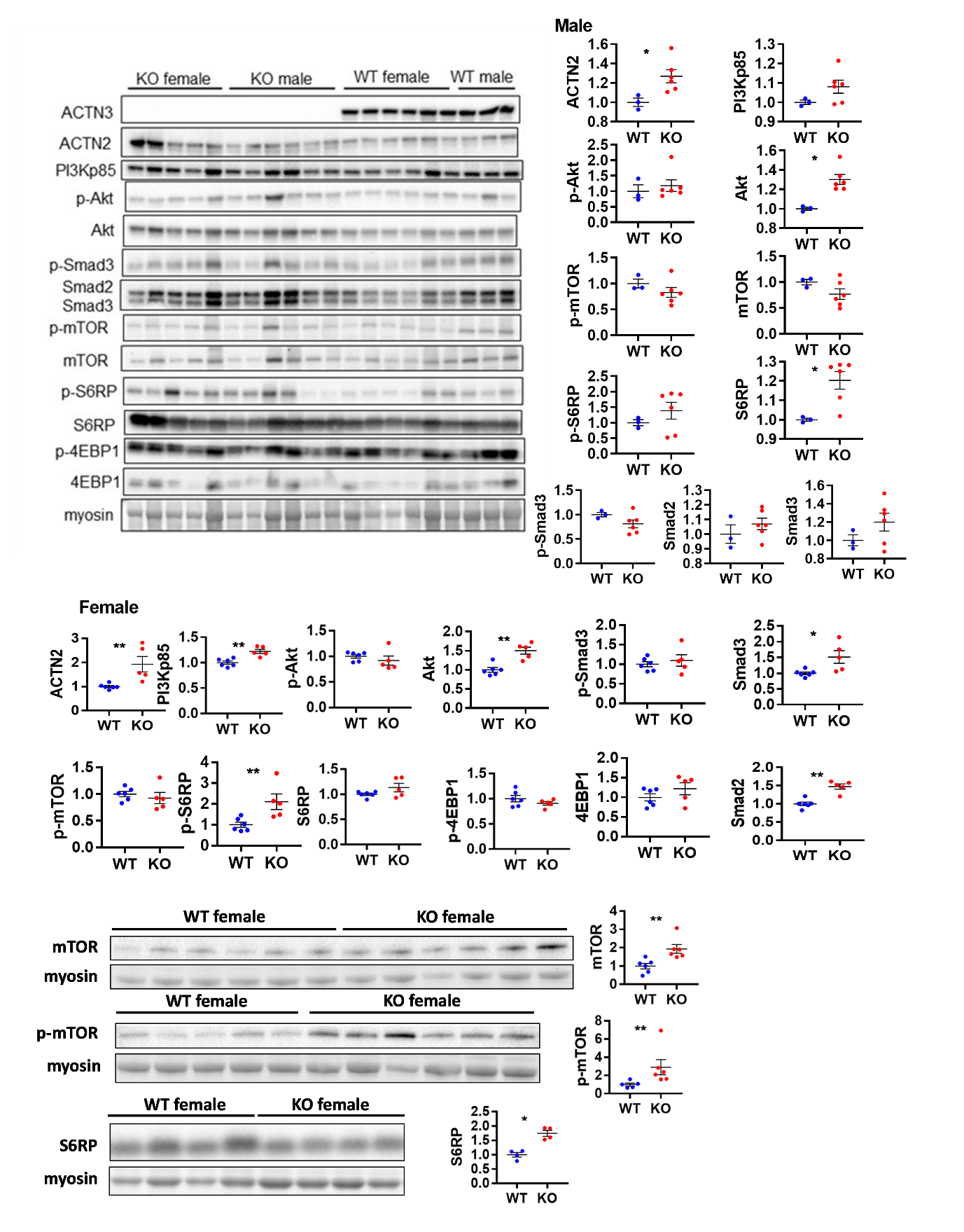

**Supplementary Figure S3**. At P28, male *Actn3* KO muscles showed significant increases in ACTN2, Akt and S6RP. Female *Actn3* KO muscles showed significant increases in ACTN2, PI3Kp85, Akt, p-mTOR, mTOR, p-S6RP, Smad3 and Smad2 compared to WT. Bottom blots were repeat western blots for mTOR, p-mTOR and S6RP for female samples. **P*<0.05, ***P*<0.01, Mann Whitney U test. *N* = 4-6 for all experiments.
