## Supplementary Figure S4 for "*ACTN3* genotype influences skeletal muscle mass regulation and response to dexamethasone"

**A.**
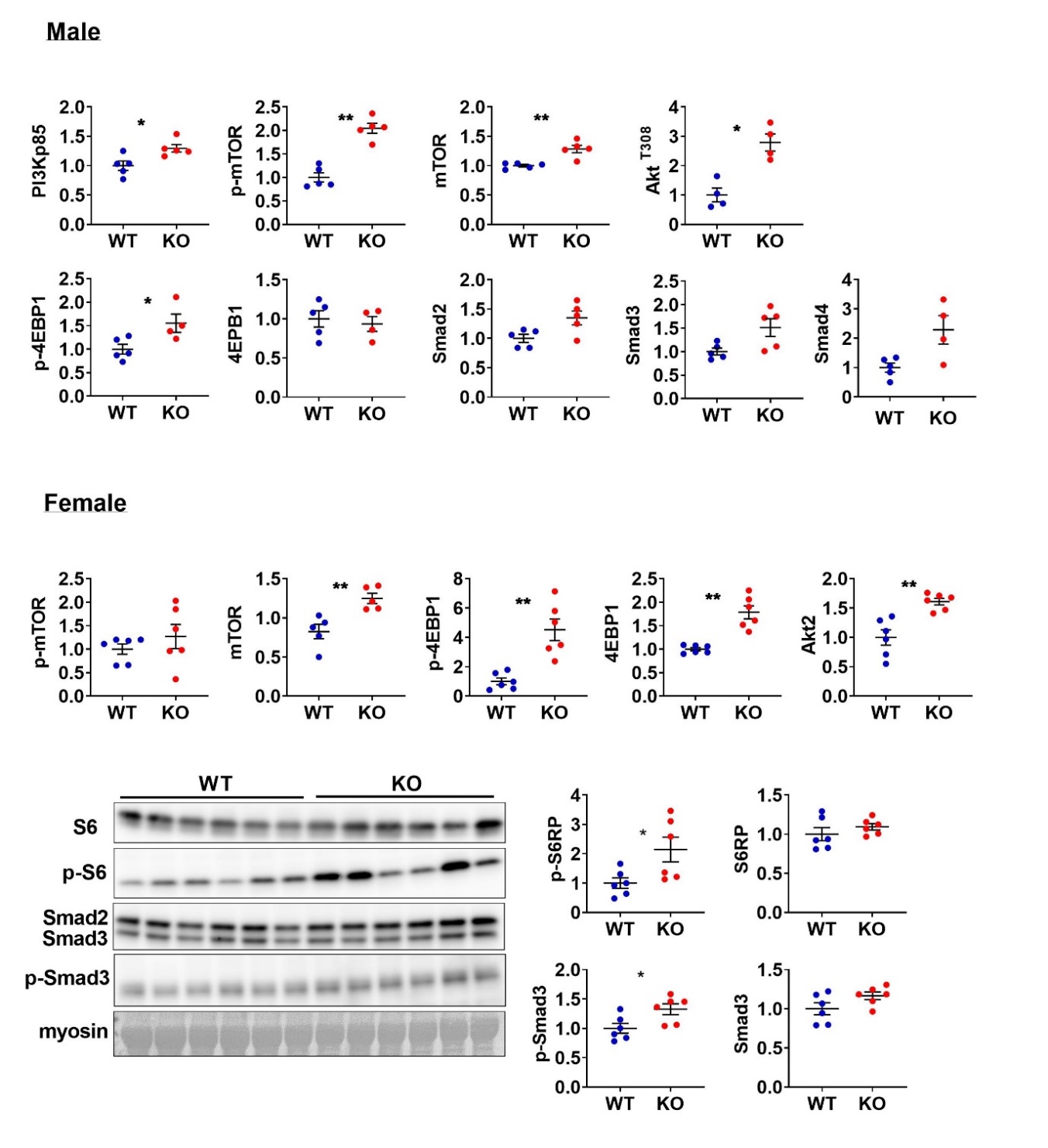

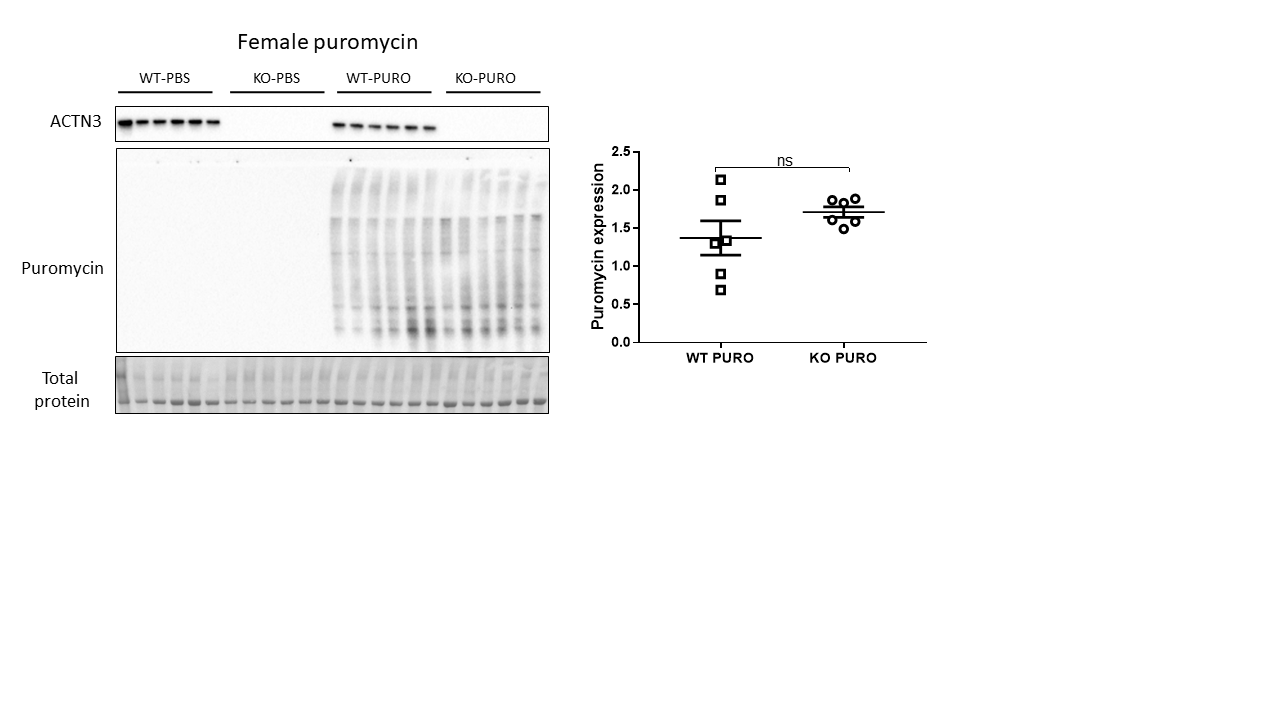
**B.**

**Supplementary Figure S4. A)** Western blot analysis of protein synthesis and breakdown signalling markers in WT and *Actn3* KO muscles at age 4 months show similar trends in male and female mice. *Actn3* KO muscles show increased activation of mTOR, 4ebp1, S6RP and Smad3 compared to WT. **B**) Protein synthesis as indicated by puromycin incorporation was not significantly different between genotypes in female mice. **P*<0.05, ***P*<0.01, Mann Whitney U test. *N* = 4-6 for all experiments.
