## Supplementary Figure S5 for "*ACTN3* genotype influences skeletal muscle mass regulation and response to dexamethasone"

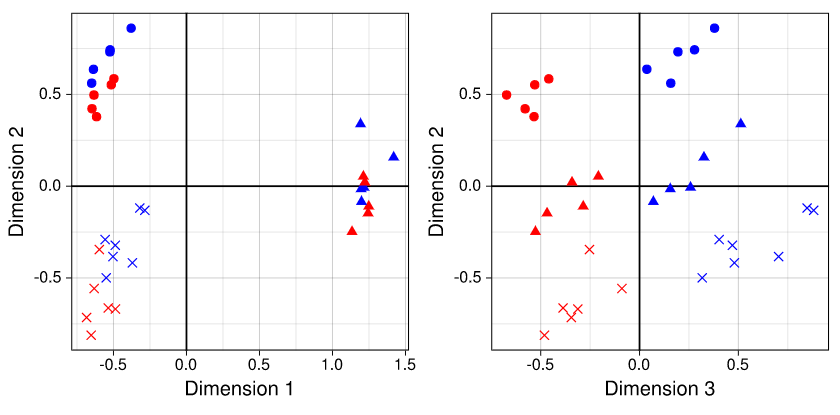

**Supplementary Figure S5.** Multidimensional scaling of gene expression levels across samples shows a broadly genotype-independent response to dexamethasone dominated by changes at t=3h post injection. Dimensions 1, 2, and 3 primarily segregate the short term drug response, a time-progressive drug response, and baseline genotype differences, respectively. Scales depict the log2 fold change of those genes which discriminate between samples.
