## Supplementary Figure S6 for "*ACTN3* genotype influences skeletal muscle mass regulation and response to dexamethasone"

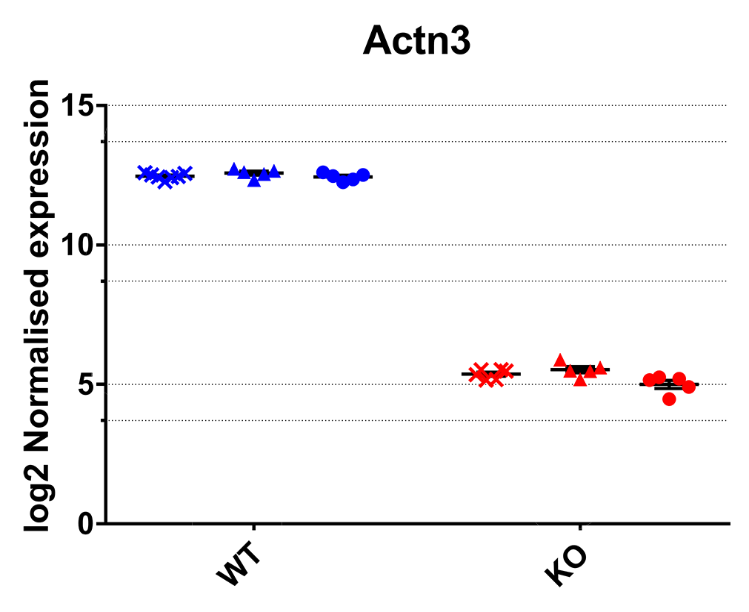

**Supplementary Figure S6.** Scatter plotting of *Actn3* expression levels confirms the efficacy of *Actn3* knockout at the transcript level in *Actn3-/-* mice. Error bars represent +/- SEM. Adjusted *p* value of differential expression with Benjamini and Hochberg’s correction =4.44x10^-37^.
