## Supplementary Figure S7 for "*ACTN3* genotype influences skeletal muscle mass regulation and response to dexamethasone"

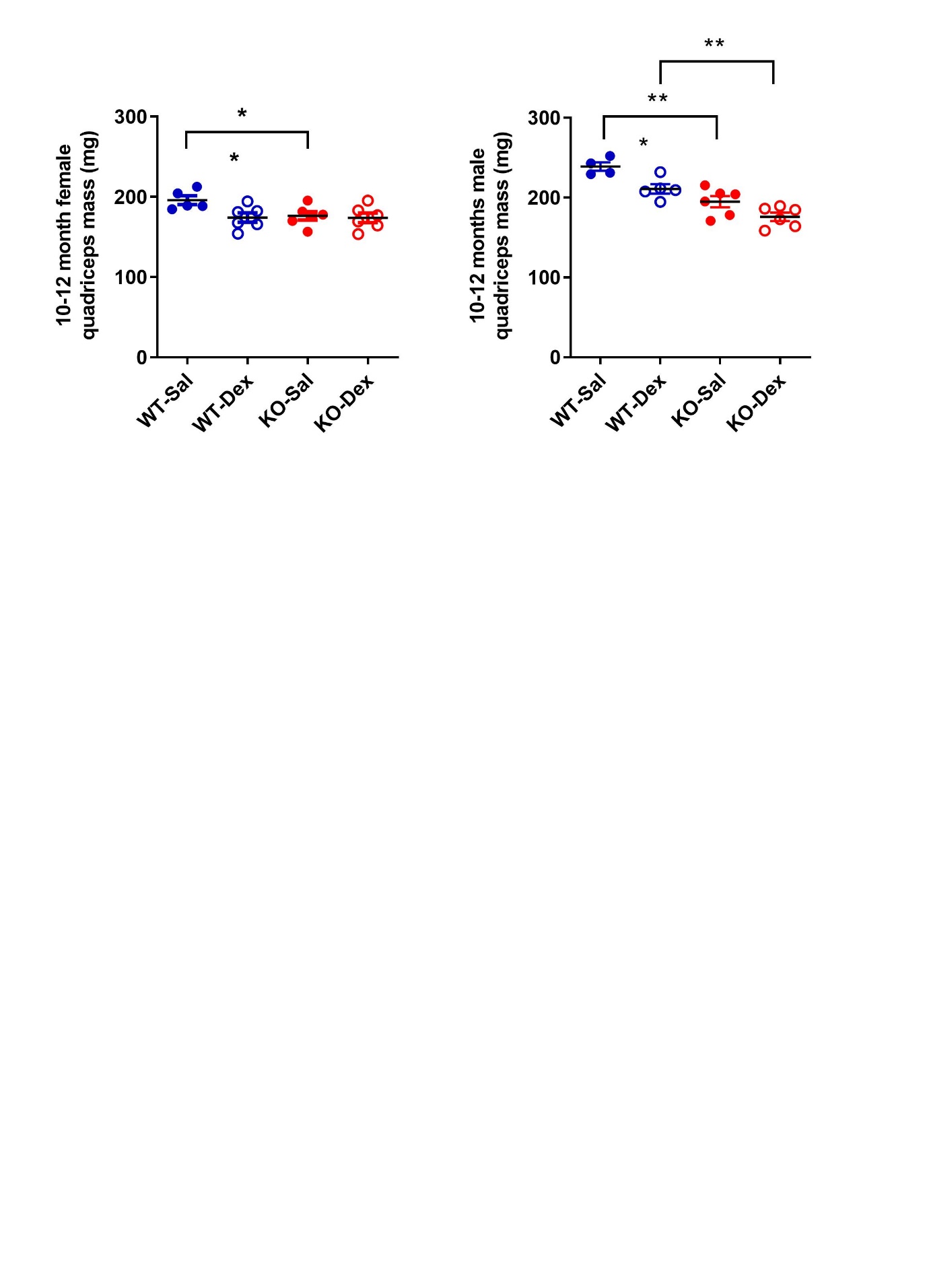

**Supplementary Figure S7.** WT male and female dexamethasone treated mice showed 11% decrease in quadriceps muscle mass relative to saline treated mice following 14 days of daily injections. Male *Actn3* KO mice also showed 9.7% lower muscle mass compared to male KO-Sal mice, however, female *Actn3* KO-Dex mice only showed 1.5% reduction in quadriceps mass compared to KO-Sal. **p*<0.05, ***p*<0.01 (Mann-Whitney U test).
